# Brain preparedness: The proactive role of the cortisol awakening response

**DOI:** 10.1101/2020.10.17.343442

**Authors:** Bingsen Xiong, Changming Chen, Yanqiu Tian, Shouwen Zhang, Chao Liu, Tanya M. Evans, Guillén Fernández, Jianhui Wu, Shaozheng Qin

**Author notes:** Equal contributions (Changming Chen, Yanqiu Tian, Shouwen Zhang, Chao Liu). Correspondence to: Shaozheng Qin, Ph.D. **Author contributions:** S.Q. conceived the experiments. B.X., C.C., Y.T., S.Z., C.L. & J.W. performed the study and data analysis. B.X., C.C., T.E., G.F. & S.Q. wrote the manuscript.

## Abstract

Upon awakening from nighttime sleep, the stress hormone cortisol exhibits a burst in the morning within 30-minutes in humans. This cortisol awakening response (CAR) is thought to prepare the brain for upcoming challenges. Yet, the neurobiological mechanisms underlying the CAR-mediated ‘preparation’ function remains unknown. Using blood-oxygen-level-dependent functional magnetic resonance imaging (BOLD-fMRI) with a dedicated prospective design and pharmacological manipulation, we investigated this proactive mechanism in humans across two fMRI studies. In Study 1, we found that a robust CAR was predictive of less hippocampal and prefrontal activity, though enhanced functional coupling between those regions and facilitated working memory performance, during a demanding task later in the afternoon. These results implicate the CAR in proactively promoting brain preparedness based on improved neural efficiency. To address the causality of this proactive effect, we conducted a second study (Study 2) in which we suppressed the CAR with a double blind, placebo controlled, randomized design using *Dexamethasone*. We found that pharmacological suppression of CAR mirrored the proactive effects from Study 1. Dynamic causal modeling analyses further revealed a reduction of prefrontal top-down modulation over hippocampal activity when performing a cognitively demanding task in the afternoon. These findings establish a causal link between the CAR and its proactive role in optimizing brain functional networks involved in neuroendocrine control and memory.

## Introduction

Upon awakening from a night sleep, cortisol, the major glucocorticoid stress hormone in humans, exhibits a burst typically by 50-160% within 30-minutes – that is known as the cortisol awakening response (CAR)(1, 2). Since its first discovery, the CAR, a hallmark of the hypothalamus-pituitary-adrenal (HPA) axis activity as well as a crucial point of reference within the healthy cortisol circadian rhythm, is thought to prepare the body for anticipated challenges of the upcoming day(3–6). In support of this “preparation” hypothesis, an individual’s CAR predicts anticipated workload, cognition and emotion(7–10), while abnormal CAR is often linked to stress-related psychopathology such as anxiety and depression(4, 11–13). However, our understanding of CAR’s neurobiological mechanisms is still in its infancy.

Cortisol acts as one of the key modulators of the human brain and cognition. It is released mainly by the zona fasciculata of the adrenal cortex in the adrenal gland(14) and can cross the blood-brain barrier to affect the neuronal excitability and functional organization of brain networks, thereby fostering behavioral adaptation to cognitive and environmental challenges(15). The conventional neurobiological models posit that glucocorticoids exert both rapid nongenomic and slower genomic actions on the limbic-frontal networks especially the hippocampus and prefrontal cortex (PFC), via high-affinity mineralocorticoid receptors (MRs) and low-affinity glucocorticoid receptors (GRs) that are co-expressed abundantly in these brain regions(16, 17). Specifically, the MR initiates rapid changes in the assembly of specific neural circuits allowing a quick and adequate response to the ongoing stressful event(18). As this process is energetically costly and may have deleterious consequences when over-engaged, MR-mediated rapid actions are complemented by slower actions via GRs on preventing these initial defence reactions from overshooting and becoming damaging. Research in animal models and humans has shown that the GR-mediated slow genomic effect on neuronal activity is not expected to start earlier than approximately 90 min after cortisol administration, and often lasts for hours(19, 20). This process can promote contextualization, rationalization and memory storage of experiences, thereby priming brain circuits to be prepared for upcoming challenges in similar contexts(17, 21). Thus, it is conceivable that the CAR, with a burst of the cortisol concentration in response to awakening in the morning, may proactively affect the brain and cognition via a similar MR/GR-mediated actions of cortisol.

Additionally, the CAR exhibits unique features that differ from conventional cortisol responses, which may involve fundamentally distinct mechanisms(22). Specifically, the CAR consists of a superimposed response to awakening, not a mere continuation of pre-awakening cortisol increase within the healthy cortisol circadian rhythm(23). It is regulated by multiple neuroendocrine and psychological processes, including i) rapid attainment of consciousness followed by slow re-establishment of one’s full alertness(2), ii) activation of hippocampal-dependent prospective memory representations for upcoming stress(4), and iii) an interplay with concurrent catecholaminergic activation when facing demanding tasks(24). Moreover, findings from previous studies point to a critical role of hippocampal and prefrontal involvement in regulating CAR(4). Patients with lesions to the hippocampus(25) or retrograde amnesia(26), for instance, do not exhibit a reliable CAR. Its magnitude also negatively correlates with prefrontal cortical thickness(27), suggesting prefrontal involvement in the CAR. In addition, functional organization of hippocampal-prefrontal networks is crucial for regulating information exchange and flexible reallocation of neural resources in support of higher-order cognitive processing such as executive function and memory(28, 29). Little, however, is known regarding the neurobiological mechanisms of how the CAR-mediated specific “preparation” function proactively modulates the human brain for higher-order cognitive functions. Based on the forementioned unique features of the CAR and empirical observations, we hypothesized that CAR would prepare the brain for upcoming demands of the day ahead via optimizing the functional organization of hippocampal and prefrontal systems.

We tested this hypothesis across two studies using blood-oxygen-level-dependent functional magnetic resonance imaging (BOLD-fMRI) with a prospective design and pharmacological manipulations dedicated to CAR (**Fig. 1a & 3a**). We opted for a well-established working memory (WM) paradigm to probe task-invoked neural activation and deactivation, especially in the dorsolateral prefrontal cortex (dlPFC) and the hippocampus, respectively(30, 31). Such functional balance between these two neurocognitive systems is known to enable a flexible reallocation of neural resources to support higher-order executive function while inhibiting task-irrelevant interference(28, 31, 32), making this domain an ideal model for studying human prefrontal-hippocampal interaction. In Study 1, 60 participants (8 of them were excluded from further analyses due to either invalid CAR data or excessive head motion during scanning; *SI Appendix, SI Methods & Table S1*) underwent fMRI while performing the WM task with low and high cognitive demands after about 6-hours relative to awakening (i.e., 14:45-15:45) in the afternoon of the same day. Six salivary samples were obtained to assess the CAR in the morning and cortisol levels before and after fMRI scanning. We observed that individuals with a robust CAR exhibited less hippocampal and prefrontal activity, as well as enhanced functional coupling between those regions and better WM performance during the cognitively demanding task.

**Fig. 1.**
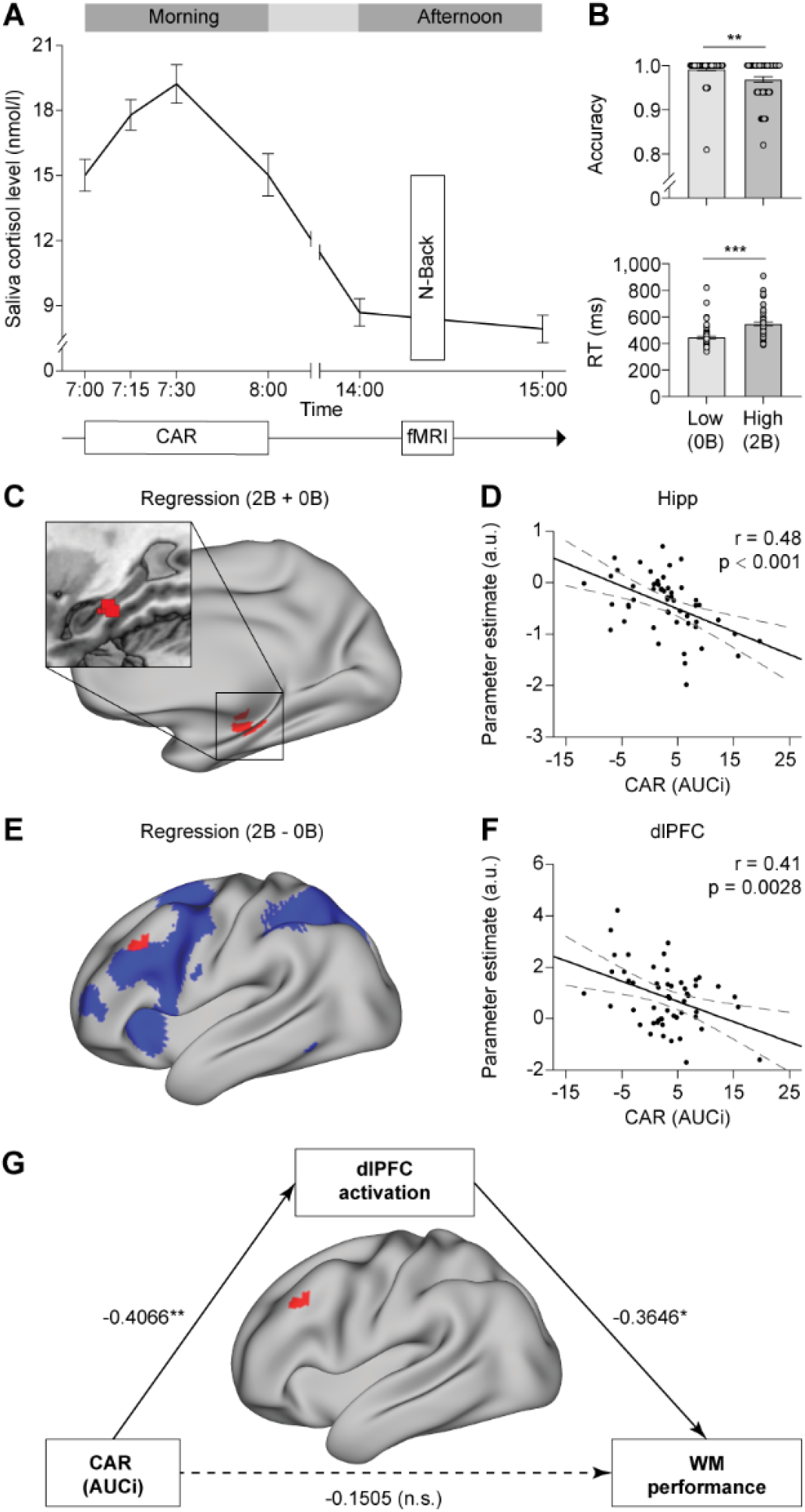
Experimental design, cortisol awakening response (CAR), and CAR-related proactive effect on brain systems from Study 1. (A) Salivary cortisol levels at 4-time points after awakening in the morning and 2-time points right before and after fMRI scanning during working-memory (WM) task about 6-hours later in the afternoon. (B) Behavioral performance on accuracy and response time (RT). (C-D) Significant cluster in the hippocampus with a negative correlation between individual’s CAR and hippocampal activity in general. (E-F) Significant cluster in the dorsolateral prefrontal (dlPFC, in red) overlapping with the main effect of WM-loads (in blue). Scatter plot depicts a negative correlation between individual’s CAR and WM-related dlPFC activity. (G) The mediating effect of the dlPFC activity on the association between the higher CAR and better WM performance. Paths are marked with standardized coefficients. Notes: Hipp, hippocampus; 0B, 0-back; 2B, 2-back; AUCi, area under the curve with respect to the cortisol increase; *, P < 0.05; **, P < 0.01; ***, P < 0.001.

To further test whether there is a causal link between an individual’s CAR and its proactive effects on task-related prefrontal and hippocampal activity, we conducted a pharmacological fMRI experiment (Study 2) by implementing a randomized, double-blind, placebo-controlled design. Sixty-three participants (4 of them were excluded from further analyses due to excessive head motion; *SI Appendix, SI Methods* & *Table S1*) received either 0.5-mg *dexamethasone* (DXM) or placebo at 20:00 on Day 1 to suppress their CAR on Day 2, allowing us to investigate its effect on task-invoked brain activity about 6-hours post-wakening. DXM, a synthetic glucocorticoid, can temporally suppress CAR via imitating negative feedback from circulating cortisol to adrenocorticotropic hormone-secreting cells of the pituitary(33, 34). Saliva samples were collected at 15-time points spanning over three consecutive days. Other procedures were similar to Study 1 (*SI Appendix, SI Methods*). As expected, we found that pharmacological suppression of CAR resembled the proactive effect from Study 1. Dynamic causal modeling further revealed a reduction of prefrontal top-down modulation over hippocampal activity during the active task 6 hours later in the afternoon. Our findings therefore establish a causal link between CAR and its proactive role in preparing hippocampal-prefrontal networks involved in WM processing.

## Results

### A robust CAR proactively predicts less hippocampal and prefrontal activity during WM

We first assessed the overall CAR profile and diurnal cortisol levels for participants from Study 1. As shown in **Fig. 1A**, cortisol levels peaked 30-minutes after awakening, followed by a decline at 60-minutes, and remained relatively low yet stable in the afternoon [*F*_5, 306_ = 36.93, *P* < 0.001]. This pattern is consistent with findings from previous studies (1, 22). To verify the effectiveness of WM-load manipulation, we conducted Separate paired t-tests on accuracy and RTs. This analysis revealed lower accuracy and slower reaction times (RTs) [both *t*_51_ > 3.43, *P* < 0.001] in the high relative to the low task demand condition (**Fig. 1B**). To identify brain systems involved in WM processing, we conducted whole-brain analyses by contrasting 2- with 0-back condition and vice versa. These analyses replicated robust WM-related activation and deactivation in widespread regions in the frontoparietal network (FPN) and default mode network (DMN) respectively (30, 31). Regions in the FPN include the dorsolateral prefrontal cortex (dlPFC) and intraparietal sulcus (IPS), and regions in the DMN include the posterior cingulate cortex (PCC), the medial prefrontal cortex and the hippocampus (*SI Appendix, Fig. S1*).

Next, we examined via whole-brain regression analyses how an individual’s CAR modulates brain functional activity involved in upcoming WM processing in the afternoon, while controlling for potential confounding factors including sleep duration, perceived stress, state and trait anxiety (*SI Appendix, SI Methods*). The area under the curve with respect to the cortisol increase (AUCi) within 1-hour after awakening was computed to quantify the overall CAR and used as the predictor of interest. We observed a hippocampal cluster [Cluster-level *P* < 0.05 family-wise error (FWE) corrected; **Fig. 1C**; *SI Appendix, Table S2]*, with lower-CAR predictive of higher hippocampal activation (or less deactivation) regardless of task demands (**Fig. 1D**). Critically, we also identified clusters in the dlPFC and the intra-parietal sulcus [Cluster-level *P* < 0.05 FWE corrected; **Fig. 1e**; *SI Appendix, Fig. S2A* & *Table S2*] with lower CAR predictive of more task-invoked prefrontal activation in the high (vs. low) demanding condition (**Fig. 1f**; *SI Appendix, Fig. S2B*). Furthermore, we found a mediating effect of the dlPFC activity on the association between the CAR and WM performance (Indirect Est. = 0.15, 95% CI = [0.026, 0.31]), indicating that robust CAR proactively promotes better WM performance via less dlPFC activation (**Fig. 1G**).

### Interaction between CAR and task demands on hippocampal and prefrontal activity

To further characterize the interaction effect between CAR and task-invoked brain activity, we conducted a set of complementary analyses by splitting participants into a robust- or lower-CAR group (defined by more than or less than 50% increase at 30-minutes after awakening, respectively) according to the criterion by previous studies(11, 35). Indeed, an independent-sample t-test confirmed a significant rise of cortisol level after awakening in the robust-relative to lower-CAR group [*t*_50_ = 8.31, *P* < 0.001] (**Fig. 2A**), but no difference in cortisol levels either before or after fMRI scanning in the afternoon [all *P* > 0.14] (**Fig. 2B**). There was no group difference in other behavioral and affective measures [all *P* > 0.66] (*SI Appendix, Fig. S3 & Table S1*). A whole-brain 2 (Group: robust- vs. lower-CAR)- by-2 (Load: low vs. high) repeated-measure analysis of variance (ANOVA) revealed a main effect of Group in the hippocampus [*F*_1,50_ = 21.54, *P* < 0.001, *η^2^* = 0.30] (**Fig. 2C&D**), an interaction effect in the dlPFC [*F*_1,50_ = 9.037, *P* = 0.004, *η^2^* = 0.15] (**Fig. 2E&F**) and the intraparietal sulcus (*SI Appendix, Fig. S4 & Table S3*) (Cluster-level *P* < 0.05 FWE corrected). Remarkably, these regions closely overlap (**Fig. 2C&E**) with those from the above-described regression analyses, highlighting the robustness of our observations. These results indicate that individuals with lower-CAR show higher hippocampal activation regardless of task demands, and higher dlPFC activation specific to a high task demand.

**Fig. 2.**
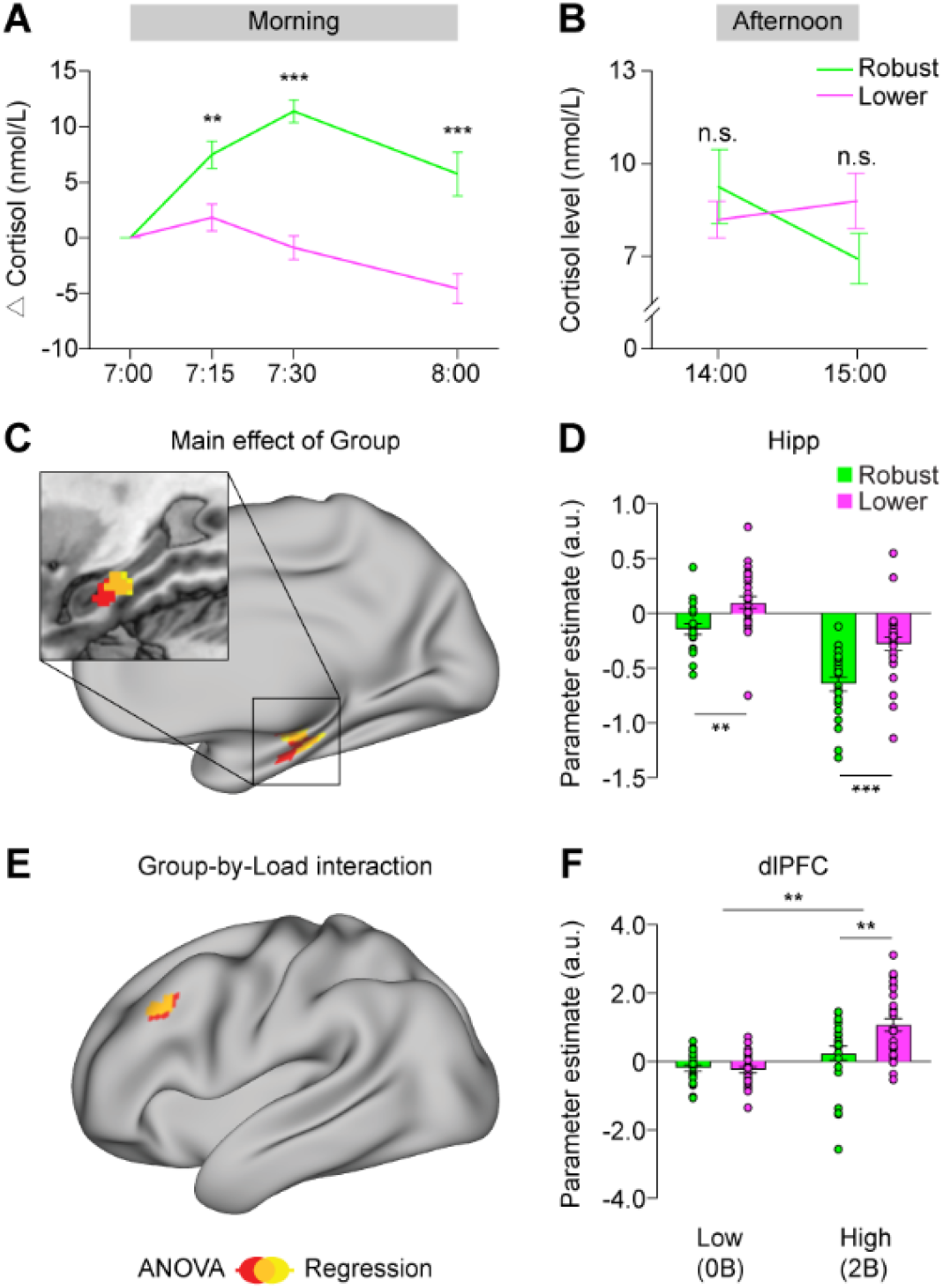
Brain systems showing higher activation in individuals with lower-than robust-CAR from Study 1. (A-B) Cortisol levels in individuals with robust- and lower-CAR in the morning, before and after fMRI scanning in the afternoon. (C) Significant clusters showing a main effect of Group in the hippocampus (in red) and overlapping (in orange) with the one (in yellow) from the regression analysis. (D) Bar graphs depict hippocampal hyper-activation regardless of WM-loads in individuals with lower-than robust-CAR. (E) Significant cluster in the dlPFC (in red) showing an interaction effect between WM-loads and Group, and overlapping (in orange) with the one (in yellow) from the regression analysis. (F) Bar graphs depict hyper-activation in the dlPFC in individuals with robust-than lower-CAR only in high (2-back) but not low (0-back) task demand. Notes are the same as Fig. 1.

### Effectiveness of pharmacological suppression of the CAR and related control measures

A pivotal question following the above-described observations is whether there is a causal link between an individual’s CAR and its proactive effects on task-related prefrontal and hippocampal activity. To address this question, we conducted a second study (Study 2) in which we suppressed the CAR with a double blind, placebo controlled, randomized design using DXM (*SI Appendix, SI Methods*). As expected, DXM administration on Day 1 suppressed participant’s CAR in the morning on Day 2, as indicated by the main effect of Group [*F*_1,57_ = 16.78, *P* < 0.001, *η^2^* = 0.23] from a 2 (Group: DXM vs. placebo)-by-15 (Time: 15-samples) ANOVA. We also observed Group-by-Time interaction effect [*F*_14,798_ = 19.91, *P* < 0.001, *η^2^* = 0.26]. Post-hoc tests revealed a flattened CAR at 0-, 15-, 30- and 60-minutes after morning awakening in DXM group [All *P* < 0.001], but no significant group differences in cortisol levels before and after fMRI scanning nor in the CAR on Day 3 when compared to placebo [All *P* > 0.18] (**Fig. 3A**). There was no significant group difference either in subjective mood across the 15-time points over three consecutive days (*SI Appendix, Fig. 5SA*), behavioral performance, sleep duration, perceived stress nor anxiety (*SI Appendix, Fig. S5B&C & Table S1*) [All *P* > 0.18]. The effectiveness of the WM-load manipulation was evidenced by separate repeated-measure ANOVA for both accuracy [*F*_2,114_ = 7.58, *P* < 0.001] and RTs [*F*_2,114_ = 48.67, *P* < 0.001] during WM. Thus, as intended, the DXM administration selectively suppressed the CAR of the experimental day, but it did not alter cortisol levels before and after fMRI scanning nor affective measures over three days.

**Fig. 3.**
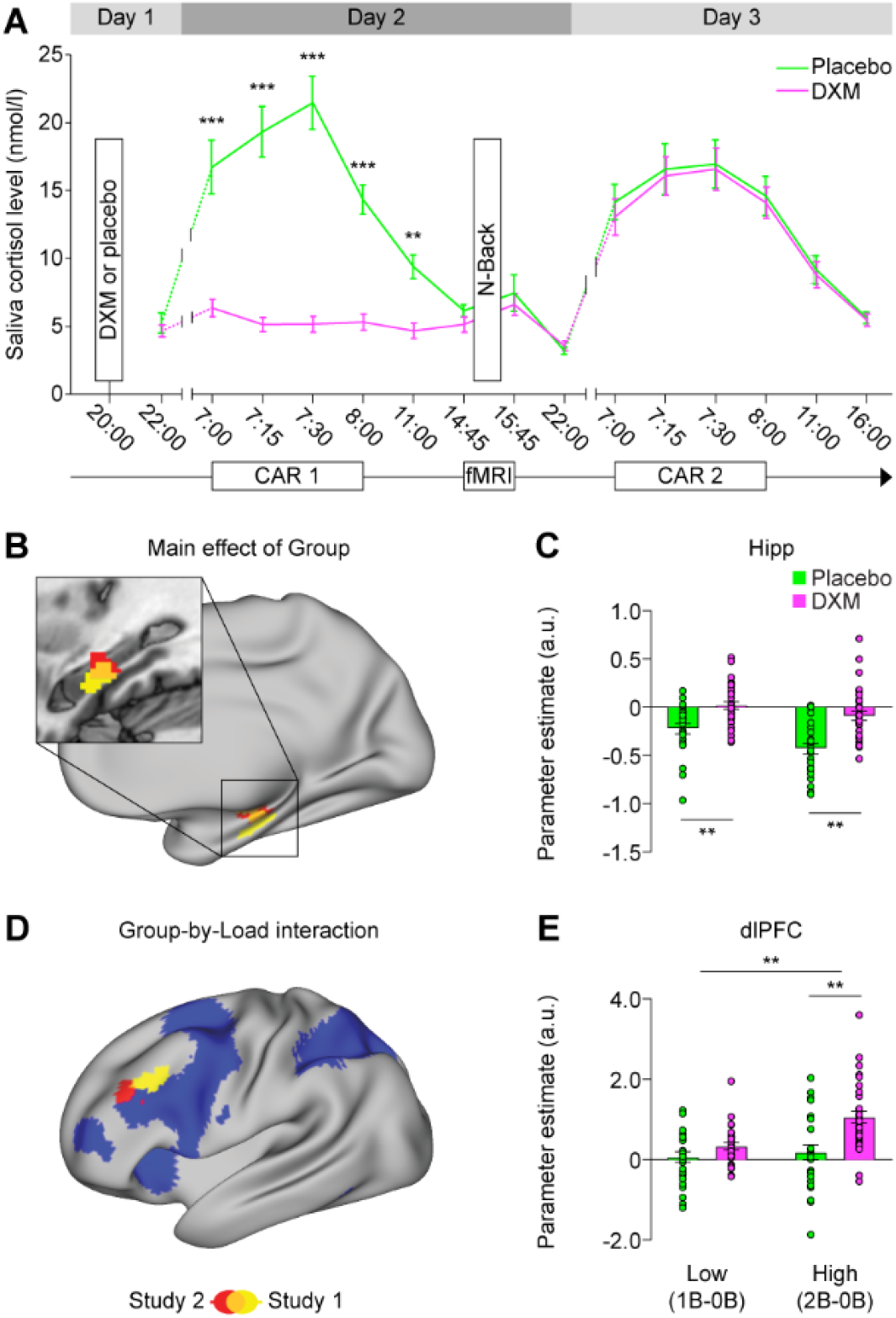
Experimental design, pharmacological suppression of CAR and its effects on brain systems from Study 2. (A) Salivary cortisol levels at 15-time points through three consecutive days. Participants received 0.5-mg either Dexamethasone (DXM) or placebo at 22:00 before sleep in the evening on Day 1. The CAR measured on Day 2 and Day 3, and fMRI data were acquired while performing a WM task with 0-, 1- and 2-back conditions in the afternoon on Day 2. (B-C) Significant cluster in the hippocampus (in red) showing general hyper-activation in DXM (vs. placebo) group which is overlapped (in orange) with the one observed in individuals with lower- vs. robust-CAR from Study 1 (in yellow). (D-E) Significant cluster in the dlPFC (in red) showing hyper-activation in the left dlPFC in DXM (vs. placebo) group only during high (but not low) task demand. Clusters in blue represent WM-related brain activation, and the cluster in green shows Group-by-Load interaction from Study 1. Notes are the same as Fig. 1.

### Suppressed CAR proactively leads to an increase in hippocampal and prefrontal activity

We then investigated whether CAR suppression using DXM in Study 2 could resemble our observed prefrontal and hippocampal hyper-activation above in individuals with lower-CAR from Study 1. We conducted a whole-brain 2 (Group: DXM vs. placebo)-by-2 (Load: Low vs. High) ANOVA. This analysis revealed a main effect of Group in the hippocampus (Cluster-level *P* < 0.05 FWE corrected, **Fig. 3B**; *SI Appendix, Table S4*) and a Group-by-Load interaction effect in the dlPFC (Cluster-level *P* < 0.05 FWE corrected, **Fig. 3D**; *SI Appendix, Table S4*). As shown in **Fig. 3B&D**, these two regions overlapped closely with the findings of Study 1, with a general hippocampal hyper-activation regardless of cognitive load [*F*_1,57_ = 26.95, *P* < 0.001, *η^2^* = 0.32] (**Fig. 3C**) and a prefrontal hyper-activation specific to high (vs. low) WM-load in the DXM as compared to the placebo group [*F*_1,57_ = 12.95, P < 0.001 *η^2^* = 0.19] (**Fig. 3E**). Other clusters are shown in *SI Appendix, Fig. S6* (*Table S4*). Thus, results from Studies 1 and 2 converge onto a causal link between the CAR and its proactive effects on task-invoked activity in the dlPFC and hippocampus about 6 hours later in the afternoon of the same day.

### Suppressed CAR reduces prefrontal-hippocampal functional coupling during WM

The above localization of brain activation linked to the CAR, however, provides limited insight into how cortisol hours later affects nuanced coordination of brain networks to support human WM. To test for CAR-mediated effects on prefrontal network properties, we implemented a generalized form of psychophysiological interaction (gPPI) analysis(36) to assess task-dependent functional connectivity of a specific seed (the dlPFC here; **Fig. 4A**) to the rest of the brain in Study 1 and 2. The dlPFC-seeded connectivity maps were then submitted to a 2 (Group)-by-2 (Load) ANOVA for statistical testing. This analysis revealed a Group-by-Load interaction in the hippocampus in Studies 1 and 2 independently (**Fig. 4B**; Cluster-level *P* < 0.05 FWE corrected; *SI Appendix, Table S5*), with weaker dlPFC-hippocampal connectivity in individuals with lower- (or DXM-suppressed) CAR as compared to those with robust-CAR (or placebo), under high but not low task demands [Study 1: *F*_1,55_ = 6.64, *P* = 0.013, *η^2^* = 0.12; Study 2: *F*_1,57_ = 23.77, *P* < 0.001, *η^2^* = 0.29] (**Fig. 4C&D**; *SI Appendix, Fig. S7*). Notably, analyses of dlPFC-hippocampal intrinsic functional connectivity at resting state showed no group difference in the two studies [All *P* > 0.47]. These results suggest that lower-/DXM-suppressed CAR proactively reduces prefrontal-hippocampal coupling during a cognitively demanding state.

**Fig. 4.**
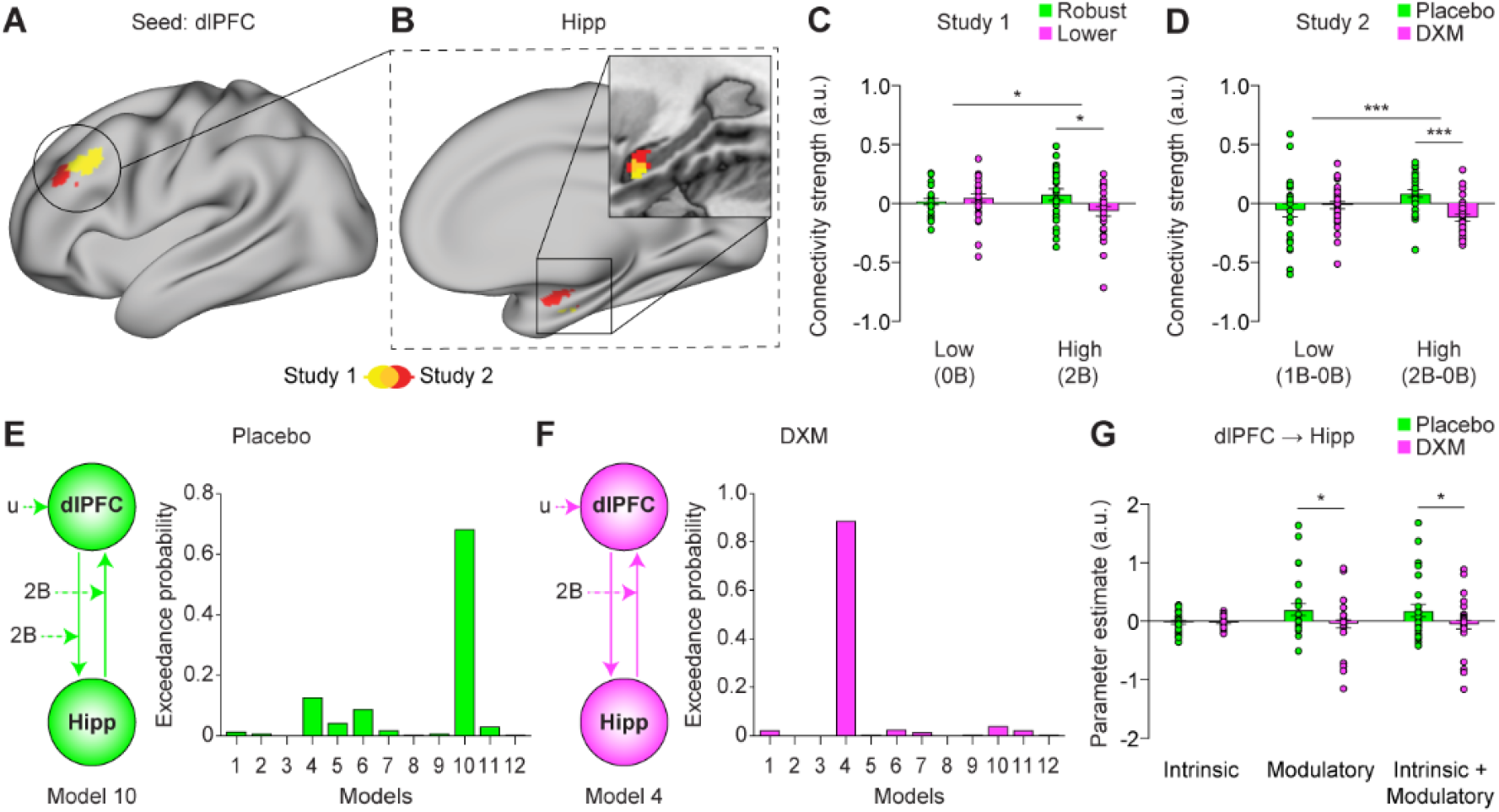
Proactive effects of the CAR on prefrontal-hippocampal dynamic functional interactions. (A) The dlPFC serving as the seed for task-dependent functional connectivity analysis. (B) Significant clusters in the hippocampus showing Group-by-Load interaction effect in Study 1 and 2. (C-D) Bar graphs depict weaker dlPFC functional coupling with the hippocampus in robust- (or placebo) than lower-CAR (or DXM) group during high but low task demand. (E) Model evidence in placebo group from dynamic causal modeling analysis favored the 10th model: inputs to the dlPFC drives the network, and high task demand (i.e., 2-back) modulates dynamic influences between the dlPFC and hippocampus bidirectionally. (F) Model evidence in DXM group favored the 4th model: inputs to dlPFC drives the network, while high task demand only modulates dynamic influence from the hippocampus to dlPFC. (G) Bar graphs depict greater dynamic modulation as well as greater intrinsic plus modulatory dynamic influence from the dlPFC to hippocampus in placebo than DXM group. Notes: u, driving input; Others are the same as Fig. 1.

### Suppressed CAR reduces prefrontal top-down modulation over the hippocampus

To further test the directionality of prefrontal-hippocampal connectivity, we modeled dynamic functional interactions between these two regions described above, by implementing Dynamic Causal Modeling to assess neural dynamics exerting from one region to the other(37). Bayesian model selection was used to identify the optimal model structure of 36 variants (*SI Appendix, SI Methods, Fig. S8&9*) that accounts best for the data in each group. For the placebo group, model evidence based on exceedance probabilities (EP) favored a model (10^th^ variant, EP = 0.68; **Fig. 4E**) where inputs to the dlPFC drive the network, and high cognitive demand (i.e., 2-back) modulates the effective connectivity between dlPFC and hippocampus bidirectionally. Model evidence for DXM, however, favored a model (4^th^ variant, EP = 0.89; **Fig. 4F**) in which inputs to the dlPFC also drives the network, but high demand only modulates the network coupling from the hippocampus to the dlPFC. Dynamic coupling parameters from the dlPFC to the hippocampus during high demand were obtained using Bayesian model averaging across all models. Independent-sample t-tests revealed a reduction in positive modulation of effective connectivity (i.e., the modulatory; *t*_57_ = −2.07, *P* < 0.05) as well as absolute effective connectivity (i.e., the modulatory plus intrinsic effect; *t*_57_ = −2.01, *P* < 0.05), but not intrinsic coupling alone in DXM relative to placebo group (**Fig. 4G**). The placebo group exhibited dynamic top-down modulation between the dlPFC and the hippocampus, whereas DXM suppression of the CAR selectively reduced the top-down modulation from dlPFC to the hippocampus during WM processing.

## Discussion

By leveraging cognitive neuroimaging and pharmacological manipulations across two studies, we investigated the neurobiological mechanisms underlying the proactive effects of human CAR on hippocampal-prefrontal functioning. In Study 1, we found that a robust CAR was predictive of less hippocampal activation regardless of task demands and less dlPFC activation selectively in a high task demand, as well as enhanced functional coupling between those regions and better working memory performance, about 6 hours later in the afternoon of the same day. These results implicate the CAR in proactively promoting brain preparedness based on improved neural efficiency. Critically, pharmacological suppression of CAR (Study 2) resembled this proactive effect from Study 1, indicating the robustness of our findings. Further, dynamic causal modeling revealed a reduction in prefrontal top-down modulation over the hippocampus. Our findings establish a causal link between the CAR and optimized hippocampal-prefrontal functional organization, suggesting a proactive mechanism of the CAR in promoting human brain preparedness.

### CAR promotes brain preparedness via improved prefrontal and hippocampal efficiency

Our observed CAR-related proactive effects on task-invoked activity in the dlPFC and hippocampus concur with the CAR-mediated “preparation” hypothesis(3–6) and extend the theoretical framework of glucocorticoids(16, 17, 21). Specifically, our observed less dlPFC activation in individuals with robust-CAR may implicate improved neural efficiency during WM processing, given the comparable behavioral performance between robust- and lower-CAR groups. This interpretation is further supported by the mediating effect of less dlPFC activity on the association between a robust CAR and higher WM accuracy. Indeed, an increase in neuronal efficiency has been linked to relatively weaker and focal activation in certain brain region(s)(38, 39), likely by utilizing fewer neural resources(40, 41).

The dlPFC and hippocampus are known to play antagonistic roles in WM processing, with prominent activation in the dlPFC and deactivation in the hippocampus(30, 31, 42, 43). Such activation/deactivation enables a flexible reallocation of neural resources between hippocampal and prefrontal systems to support executive functions(28, 31, 32). The hippocampal deactivation, likely via a GABAergic inhibition mechanism(44), is to suppress task-irrelevant thoughts and/or mind-wondering in favor of information maintenance and updating in WM(31, 42, 43). Thus, more hippocampal deactivation (less activation) here may reflect more effective suppression of task-irrelevant thoughts in individuals with robust-than lower-CAR.

Our observation on prefrontal-hippocampal systems differs from previous findings of increased local activity in the dlPFC 4-hours after administration of exogenous corticosteroid to mimic a cortisol rise (20). Given the CAR’s unique features in morning awakening, it takes us from the general GR-mediated slow effect to a CAR-specific “preparation”. The CAR is believed to be accompanied by activation of prospective memory representations of upcoming challenges for the day ahead(4). Such mnemonic aspects of the CAR could be important determinants of its proactive effects on brain networks. According to the neurobiological models of glucocordicoids,, the brain can generate memory-dependent inhibitory traces to control cortisol responses and prime specific neural circuits to be prepared for future threats in similar contexts(17, 21), through the MR/GR-mediated actions on initiating rapid reactions, contextualizing and regulating subsequent neuroendocrinal and behavioral adaptation to stress. Mnemonic-related brain circuits, for instance, can diminish responsiveness to repeatedly exposed stimuli to save engergy consumption(21). Thus, we speculate that the CAR, via similar MR/GR-mediated actions, may proactively set up a tonic tone with memory-dependent inhibitory traces to promote neuroendocrine cotrnol and mnenomic-related brain functions, thereby improves prefrontal-hippocampal efficiency during WM processsing. Suppressed-CAR implicates a decrease in such tonic inhibitory tone, which may account for more activity in those brain circuits that were also found in individuals with lower-CAR. To take it one step further, more dlPFC activity observed only under high but not low task demands in individuals with lower/suppressed-CAR may result from an interplay between reduced tonic inhibition in the background and task-induced phasic catecholaminergic actions on prefrontal networks during WM processing(24). Most likely, this proactive effect of the CAR on improved prefrontal-hippocampal efficiency can in turn optimize a flexible reallocation of neurocognitive resources among these systems to meet ever-changing cognitive demands. Our findings below from connectivity and dynamic causal modeling further support this interpretation.

### CAR promotes brain preparedness via optimizing prefrontal-hippocampal network coupling and dynamic interactions

Beyond regional activation, a robust CAR proactively enhances functional coupling between the dlPFC and the hippocampus during WM, with higher connectivity in individuals with robust-than lower-CAR. Pharmacological suppression of CAR in Study 2 resembles this observation again. Prefrontal-hippocampal functional organization is recognized to play a critical role in various cognitive tasks including WM(45–47), through both direct and indirect neuronal connections(48, 49). Higher dlPFC coupling with the hippocampus in individuals with robust-CAR may reflect more efficient functional communication to support a flexible reallocation of neurocognitive resources to meet cognitive demands. Notably, weaker prefrontal-hippocampal coupling in individuals with lower-CAR came along with stronger dlPFC activation during high task demands. A stronger activation in the dlPFC may implicate compensation for suboptimal prefrontal-hippocampal functional organization(38, 41).

Our dynamic causal modeling further revealed that pharmacological suppression of CAR reduces the effective connectivity from the dlPFC to the hippocampus during WM about 6 hours later. Such metric has been linked to the directionality of neural dynamics that one neuronal system exerts over another(37). Thus, our observation is most likely to reflect a reduction in prefrontal down-regulation over hippocampal activity during WM. Findings from previous studies have suggested that similar down-regulation involves a goal-directed signal that originates in the dlPFC and spreads downstream via polysynaptic pathways to the hippocampus, thus integrating these regions in a task-dependent manner(50, 51). If the CAR is responsible to promotes the route of goal-directed input from the dlPFC to suppress hippocampal processing of task-irrelevant thoughts during WM, DXM-suppressed CAR in the morning would mute the dlPFC influence on this network dynamics. Indeed, two aspects of our results support this assumption. First, the placebo group favored a model with inputs to the dlPFC driving the network and high task demands modulating connectivity between the hippocampus and the dlPFC bidirectionally, whereas the DXM-suppressed CAR group favored a model with the same inputs to the dlPFC driving the network, but reduced top-down modulation of network dynamics from the dlPFC to the hippocampus during high task demands. Second, this top-down modulation showed a strong trend to be positive, i.e., according to dynamic causal modeling reduced dlPFC recruitment caused reduced hippocampal activation during WM processing.

Taken together, our findings from activation, connectivity and dynamic causal modeling converge into a model of how the CAR prepares brain networks for the upcoming challenges: the CAR-mediated tonic inhibitory tone may work in concert with task-induced phasic catecholaminergic actions, thereby proactively improving neural efficiency in hippocampal-prefrontal networks and optimizing the flexible reallocation of neurocognitive resources in these networks to support neuroendocrinal control, executive function and memory. Indeed, many accounts regarded the interplay of glucocorticoid and catecholaminergic actions on modulating not just neural activities of different systems but also the dynamic organization of large-scale brain networks(24, 52). Future studies are required to address the complex interplay of the CAR and other neuromodulatory systems.

**In conclusion**, our findings establish a causal link between the CAR and its proactive role in the functional coordination of prefrontal-hippocampal networks involved in executive functioning. Combining cognitive neuroimaging with pharmacological manipulation advances our understanding of the CAR-mediated neuromodulatory pathways for upcoming cognitive and environmental challenges, and highlights brain preparedness for the day ahead after awakening more broadly. Our study also characterizes the proactive role of CAR on brain preparedness for the day ahead after awakening and could lead to the development of useful biomarkers in both healthy and clinical populations.

## Materials and Methods

### Participants

A total of 123 young, healthy, male college students participated in two separate studies, with 60 (mean age: 21.6 ± 0.76 years old; ranged: 20 - 24 years old) in Study 1 and 63 (mean age: 22.9 ± 1.9; ranged from 18 to 27 years old) in Study 2 (see *SI Appendix, SI Methods* for details). Only men were included because of hormonal fluctuations across the menstrual cycle and the impact of hormonal contraceptives in young adult females(53). Participants reported no history of neurological, psychiatric or endocrinal disorders. Exclusion criteria included current medication treatment that affects central nervous or endocrine systems, daily tobacco or alcohol use, irregular sleep/wake rhythm, intense daily physical exercise, abnormal hearing or (uncorrected) vision, predominant left-handedness, current periodontitis, stressful experience or major life events. Informed written consent was obtained from all participants before the experiment, and the study protocol was approved by the Institutional Review Board for Human Subjects at Beijing Normal University. The protocol with pharmacological manipulation was registered as a clinical trial before the experiment (https://register.clinicaltrials.gov/; Protocol ID: ICBIR_A_0098_002).

### General Experimental Procedure

In Study 1, we explored the relationship between CAR and the neurocognitive correlates of working memory (WM) in the natural setting. Salivary samples were obtained at 6 time points to assess CAR and diurnal rhythms of cortisol levels. The brain imaging data were acquired when participants performed a numerical N-back task with two loading conditions (i.e., 0- and 2-back) in the afternoon of the same day (**Fig. 1a**).

In Study 2, we implemented a randomized, double-blinded, placebo-controlled design to investigate the causal link of CAR with brain activity during WM task. Participants orally receive either a dose of 0.5-mg Dexamethasone (i.e., DXM group) or an equal amount Vitamin C (i.e., placebo group) pill at 20:00 on Day 1. Participants completed a similar numerical N-back task with three conditions (0-, 1- and 2-back) during fMRI scanning in the afternoon on Day 2. A total of 15 saliva samples were collected through 3 consecutive days, while participant’s subjective mood was monitored concurrently by the positive and negative affection scale (PANAS)(54). The other experimental settings are identical to those of Study 1 (**Fig. 3a**).

### Physiological and Psychological Measures

Details of salivary cortisol measure, cognitive task and questionnaires are provided in *SI Methods*.

### Brain Imaging Data Acquisition

Functional brain images were collected during the N-back task using a gradient-recalled echo planar imaging (GR-EPI) sequence. High-resolution anatomical images were acquired in the sagittal orientation using a T1-weighted 3D magnetization-prepared rapid gradient echo sequence (see *SI Appendix, SI Methods* for details).

### Brain Imaging Data Analysis

#### Preprocessing

Image preprocessing and statistical analysis of fMRI data were performed using Statistical Parametric Mapping (SPM12, http://www.fil.ion.ucl.ac.uk/spm). Details of fMRI preprocessing are provided in *SI Methods*.

#### Univariate GLM analysis

To assess neural activity associated with the experimental conditions, each condition was modeled separately as boxcar regressor and convolved with the canonical hemodynamic response function (HRF) built in SPM12. The 6 parameters for head movement were also included in the model as covariates to account for movement-related variability. A high-pass filtering cutoff of 1/128 Hz and a serial correlation correction by a first-order autoregressive model (AR) were also applied. Contrast images for each condition, generated at the individual level fixed-effects analyses, were submitted to a second-level group analysis treating participants as a random factor (see *SI Appendix, SI Methods* for details).

#### Structural equation modeling

Structural equation models (SEMs) were constructed to examine the hypothesized mediating effects of prefrontal activation on the associations between the CAR and WM performance using Mplus 7.0 (see *SI Appendix, SI Methods* for details).

#### Task-dependent functional connectivity analysis

To examine whether the hyper-activation caused by suppressed CAR was related to dlPFC coupling with brain regions, we conducted generalized psychophysiological interaction (gPPI) analysis(36) (see *SI Appendix, SI Methods* for details).

#### Dynamic causal modeling

To further investigate how suppressed CAR modulates functional interactions between the dlPFC and the hippocampus (ROIs identified from the above activation analysis) during WM, we estimated the effective connectivity between these two brain regions using dynamic causal modeling (DCM)(37) (see *SI Appendix, SI Methods* for details).

## Acknowledgments

We thank W. Lin, L. Zhuang and L. Hao for their assistance in data collection and data analysis.

## Funding

This work was supported by the National Natural Science Foundation of China (31522028, 81571056, 31871110).

## Data and Code Availability

The codes that support findings of this study are available from https://github.com/QinBrainLab/2020_CAR_preparedness. The original data are available from the corresponding author pending on reasonable request.

## Notes

**Competing interests:** The authors declare no competing interests.

### Competing Interest Statement

The authors have declared no competing interest.

