## Supplemental Information for "Brain preparedness: The proactive role of the cortisol awakening response"

Shaozheng Qin

**This PDF file includes:**

Supplementary text
Figures S1 to S9
Tables S1 to S5
SI References

**Supplementary Information Text**
**Materials and Methods**

Participants

*Exclusions.* Data from 12 participants were excluded from the analyses due to excessive (beyond 2 mm/degree) head movement during scanning (5 and 3 participants for Study 1 and 2, respectively) or incomplete salivary samples (3 and 1 participants for Study 1 and 2, respectively). Thus, a final sample of 52 participants (28 and 24 participants for robust- and lower-CAR groups, respectively) were included in Study 1, and another sample of 59 participants (26 and 33 for placebo and DXM groups, respectively) were included in Study 2 (*SI Appendix, Table S1*).

Physiological and Psychological Measures

*Salivary cortisol measure.* Cortisol levels were measured from saliva samples. In Study 1, a total of 6 saliva samples were collected, with 4 time points spanning within 1 hour immediately after awakening (i.e., 0, 15, 30 and 60 minutes) and 2 time points right before and after fMRI scanning (**Fig. 1a**). In Study 2, participants orally took a pill of 0.5-mg DXM (or placebo) at 20:00 on Day 1, then a total of 15 saliva samples (i.e., S0 to S14) were collected to cover cortisol's diurnal rhythms across 3 consecutive days at the following time points: on Day 1 at 22:00; on Day 2 at 0 min (7:00), 15 min (7:15), 30 min (7:30) and 60 min (8:00) after awakening ; 11:00 before lunch, right before and after the fMRI scanning in the afternoon from 14:00 to 17:00, and 22:00 in the evening; on Day 3 at 0 min (7:00), 15 min (7:15), 30 min (7:30) and 60 min (8:00) after awakening, 11:00 and 16:00 respectively(**Fig. 3a**). Saliva was collected using Salivette collection device (Sarstedt, Germany). Participants were asked not to brush their teeth, drink or eat within 1 hours before sampling in order to avoid saliva contamination. They were also required to refrain from any alcohol, coffee, nicotine consumption as well as excessive exercise at least one day before the experiment. Several steps were adopted to increase compliance and the quality of saliva collection for CAR. First, participants were explained the importance of adhering to the sampling protocol and were required to follow the instructions carefully. Each participant was given a copy of the point-by-point instruction sheet and informed by a phone call before sleep to increase compliance for upcoming morning sampling(1, 2). Second, they were instructed to set alarms to remind them the sampling timing of each sampling. The wake-up alarm was set around 7:00 am. Third, to objectively monitor participants' sleep and salivary sampling, a motion watch (MW8; CamNtech, UK) was used to record their movement at night and in the morning which helped identify the sleep quality and waking up time, a medication event monitoring system (MEMS; AARDEX, Switzerland) was also used to track the exact time of each sample collection.

Salivary samples were returned back to the laboratory and kept frozen (−20 °C) until the assay. After thawing and centrifuging at 3000 rpm for 5 min, the samples were analyzed using an electrochemiluminescence immunoassay (ECLIA, Cobas e601, Roche Diagnostics, Mannheim, Germany) with sensitivity of 0.500 nmol/L (lower limit) and a standard range in assay of 0.5–1750 nmol/L. Intra and inter-assay variations were below 10%. The CAR was computed by the area under the curve with respect to increase (AUCi) by the following equation: $AUCi = (S1+ S2) \times 0.25/2 + (S2 + S3) \times 0.25/2 + (S3 + S4) \times 0.5/2 - S1 \times (0.25 + 0.25 + 0.5)$. S1 to S4 represent the measurements of 4 samples collected within 1 hour immediately after awakening. The AUCi reflects the dynamics of the cortisol awakening response and emphasizes diurnal changes over time(3, 4).

*Cognitive task.* A blocked-design N-back task was used in both studies. In study 1, the entire task included 10 blocks of alternating 0- and 2-back conditions. In study 2, the task consisted of 12 blocks of alternating 0-, 1- and 2-back conditions. Each block started with a 2-s cue indicating the experimental condition, followed by a pseudo-randomized sequence consisting of 15 digits. Each digit was presented for 400 ms, followed by an inter-stimulus-interval of 1400 ms. The blocks were interleaved by a jittered fixation ranged from 8 to 12 s, resulting in a mean inter-block duration of 38 s. During the 0-back condition, participants were instructed to detect whether the current digit was '1'. During the 1-back condition, participants were instructed to detect whether the current digit had appeared 1 position back in the sequence. During the 2-back condition, participants were instructed to detect whether the current digit had appeared 2 positions back in the sequence. Each sequence contained either 2 or 3 targets, and participants were asked to make a button press with their right index finger as fast as possible when detecting a target.

*Questionnaires*. When participants arrived at the laboratory in both Study 1 and 2, they received two questionnaires: the State-Trait Anxiety Inventory (STATI)(5) measuring the participants' state and trait anxiety, and the Perceived Stress Scale (PSS)(6) measuring their long term psychological stress levels. A sleep questionnaire was also used to log participants' sleeping quality across the experimental days. In Study 2, subjective mood state was also assessed using the PANAS at time points coinciding with collection of saliva samples.

Brain Imaging Data Acquisition

Whole-brain images in both studies were acquired on a Siemens 3.0 Tesla TRIO MRI scanner (Erlangen, Germany) in the National Key Laboratory of Cognitive Neuroscience and Learning & IDG/McGovern Institute for Brain Research at Beijing Normal University. Functional brain images were collected during the N-back task using a gradient-recalled echo planar imaging (GR-EPI) sequence (axial slices = 33, volume repetition time = 2.0 s, echo time = 30 ms, flip angle = 90°, slice thickness = 4 mm, gap = 0.6 mm, field of view = 200 × 200 mm, and voxel size = 3.1 × 3.1 × 4.6 mm). High-resolution anatomical images were acquired in the sagittal orientation using a T1-weighted 3D magnetization-prepared rapid gradient echo sequence (slices = 192, volume repetition time = 2530 ms, echo time = 3.45 ms, flip angle = 7°, slice thickness = 1 mm, field of view = 256 × 256 mm, and voxel size = 1 × 1 × 1 mm3).

Brain Imaging Data Analysis

*Preprocessing.* The first four volumes of functional images were discarded for signal equilibrium and participant’s adaptation to scanning noise. Remaining images were corrected for slice acquisition timing, realigned for head motion correction, co-registered to the gray matter image segmented from the anatomical T1-weighted images, and subsequently spatially normalized into a common stereotactic Montreal Neurological Institute (MNI) space. Images were then resampled into 2-mm isotropic voxels, and finally smoothed by an isotropic three-dimensional Gaussian kernel with 6mm full-width at half-maximum. The data were statistically analyzed under the framework of general linear models (GLM).

*Univariate GLM analysis.* In Study 1, we first conducted a paired t-test to identify brain regions associated with WM by contrasting the 2- with 0-back condition and vice versa (collapsing across groups) (*SI Appendix, Fig. S1*). Given that the main effect of 2- vs. 0-back produced very strong and extensive activations, we applied a rather conservative statistical threshold to allow for a detailed characterization of peak activations ($P < 0.05$ familywise error rate (FWE) correction with Gaussian random field theory in SPM12).

To examine how individual differences in CAR modulate WM-related brain activity, we then conducted whole-brain multiple regression analysis on the contrast of 2-back plus 0-back condition (i.e., 2B+0B) and 2-back minus 0-back condition (i.e., 2B-0B), with CAR as the covariate of interest, while sleep duration, perceived stress and state-trait anxiety as covariates of no interest. Significant clusters were determined using a height threshold of $P < 0.001$ and an extent threshold of $P < 0.05$ with cluster-based FWE correction. Regions of interests (ROIs) were defined by overlapping these clusters with a template derived from automated meta-analysis of the most recent 1,091 fMRI studies with 'working memory' as a search term in Neurosynth (http://www.neurosynth.org). Correlation analyses for data extracted from these clusters were conducted and the patterns of correlation were illustrated on scatterplots.

To further characterize the interaction effect between CAR and task-invoked brain activity, we conducted a complementary analysis by splitting participants into two groups of individuals with robust- and lower-CAR. According to the criterion outlined by previous studies(7, 8), individuals whose cortisol level raised more than 50% at 30 minutes after awakening were grouped into the robust-CAR group, whereas individuals with less than 50% increase were assigned into the lower-CAR group. We first conducted independent-sample t-test to confirm the group difference of the CAR. We then conducted repeated-measure analysis of variance (ANOVA) on the whole-brain level, with WM (i.e., 0- and 2-back) as within-subject factor and Group (i.e., robust- and lower-CAR) as between-subject factor. Significant clusters were determined using a height threshold of $P < 0.001$ and an extent threshold of $P < 0.05$ with cluster-based FWE correction. The definition of ROIs was the same as above. ANOVA for data extracted from these clusters were conducted and the patterns of main effect of Group and Group-by-Load interaction effect were plotted on bar graphs.

In study 2, to test whether brain activity following suppressed CAR resembles that in Study 1, we conducted similar repeated-measure ANOVA on the whole-brain level, with WM-load (i.e., 1- and 2-back, relative to 0-back baseline) as the within-subject factor and pharmacological treatment (i.e., DXM and placebo group) as the between-subject factor. Clusters of significance were determined in the same way as in Study 1. ANOVA was conducted for data extracted from these clusters. The patterns of main effect of Group and Group-by-Load interaction effect were plotted on bar graphs.

*Structural equation modeling*. Bias corrected bootstrap was conducted (5000 samples) to test the mediating effect(9). Both direct and indirect effects of prefrontal activation on the association between individual differences in CAR and WM performance were estimated, which generated percentile based on confidence intervals (CI). All reported P values are two-tailed.

*Task-dependent functional connectivity analysis.* The dlPFC seed was defined as a cluster that showed Group-by-Load interaction from activation analysis in Study 1 and Study 2. The mean time series from the seed ROI were then deconvolved to uncover neuronal activity (i.e., physiological variable) and multiplied with the task design vector contrasting WM-load (i.e., 0-, 1- and 2-back) (i.e., psychological variable) to form a psychophysiological interaction vector. This interaction vector was convolved with a canonical HRF to form the gPPI regressor of interest. Task-related activations were also included in this GLM to remove out the effects of common driving inputs on brain connectivity. Contrast images corresponding to PPI effects at the individual level were then submitted to a group analysis. We conducted repeated-measure ANOVA on the whole-brain level, with WM-load as the within-subject factor and group as the between-subject factor. Significant clusters were determined using a height threshold of $P < 0.001$ and an extent threshold of $P < 0.05$ with cluster-based FWE correction. ANOVAs were conducted for data extracted from these clusters. The patterns of Group-by-Load interactions effect were plotted on bar graphs.

*Dynamic causal modeling.* DCM explains regional effects in terms of dynamically changing patterns of connectivity during experimentally induced contextual changes. Importantly, this method allows inferences about the direction of causal connections, i.e., whether the CAR modulates the 'top-down' connection from dlPFC to hippocampus or the reverse 'bottom-up' connection. We defined a standard model including both regions as nodes with bidirectional, intrinsic connections. This model was then modified to yield 36 models varied in the connections that could be modulated and in the locations of driving inputs during different WM-loads respectively. *SI Appendix, Fig. S8* illustrated the structures of 12 models including only 2-back as modulatory.

The models were estimated separately for each participant. To this end, we therefore extracted the regional time series of the BOLD signal for each participant. First, two ROIs were defined as clusters in the hippocampus (showing main effect of Group in activation analysis in Study 2) and the dlPFC (showing Group-by-Load interaction from activation analysis in Study 2). The first eigenvariate from a ROI adjusted for effects of interest (i.e. 0-, 1-, and 2-back) constitutes the regional activation. Model fitting was based on these data and was achieved by adjusting the model parameters to maximize the free-energy estimate of the model evidence(10).

Separated Bayesian model selection (BMS) for both DXM and placebo group was then used to identify the model that could account best for the data(11). A random-effects approach was taken, since it does not assume that the optimal model will be the best for each individual(12). This analysis reports the exceedance probability (EP), i.e., the probability to which a given model is more likely to have generated the data from a randomly selected participant than any other competing model. The group level DCM analysis was also conducted using Bayesian model averaging (BMA)(11), which is less dependent on assumptions about model structure. BMA is a Bayesian approach that averages each parameter across models (and across subjects) such that the contribution of each model (of each subject) for that parameter is weighted by the model's posterior probability. Independent-sample t-tests were conducted between groups for intrinsic coupling, modulatory, modulatory plus intrinsic effect separately.

### Supplementary Figures S1-S8

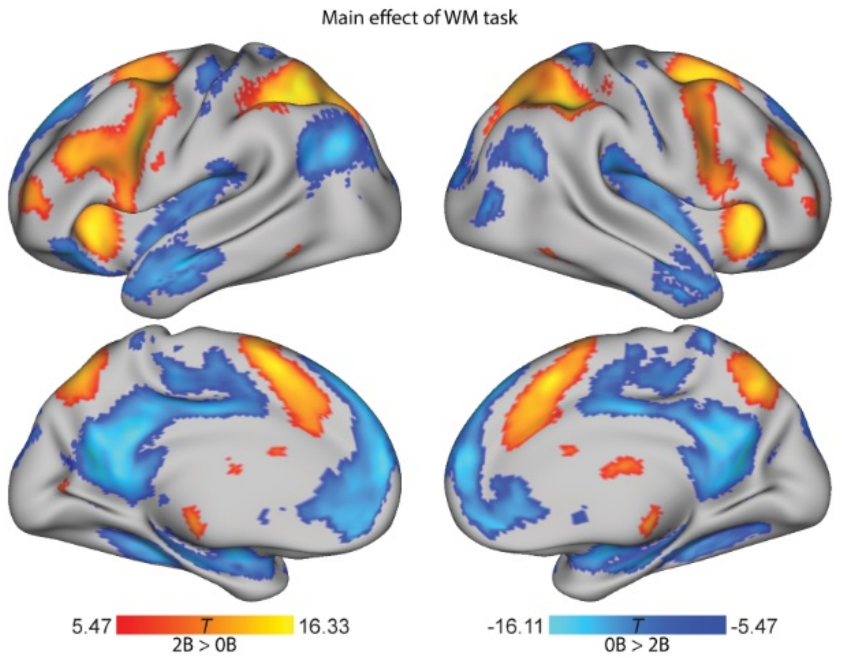


**Supplementary Fig. S1. Brain systems involved in WM processing.** Lateral and medial views of significant clusters in a widespread set of brain regions in frontoparietal executive and default mode networks. Regions in the frontoparietal executive network with positive activation includes the bilateral dorsolateral prefrontal cortex (dlPFC), intraparietal cortex (IPS) and among others. Regions in the default mode network with negative activation or deactivation include the posterior cingulate cortex (PCC), medial prefrontal cortex and the hippocampus. Statistical parametric maps are thresholded at $P < 0.05$ FWE correction. Notes: 0B, 0-back; 1B, 1-back; 2B, 2-back.

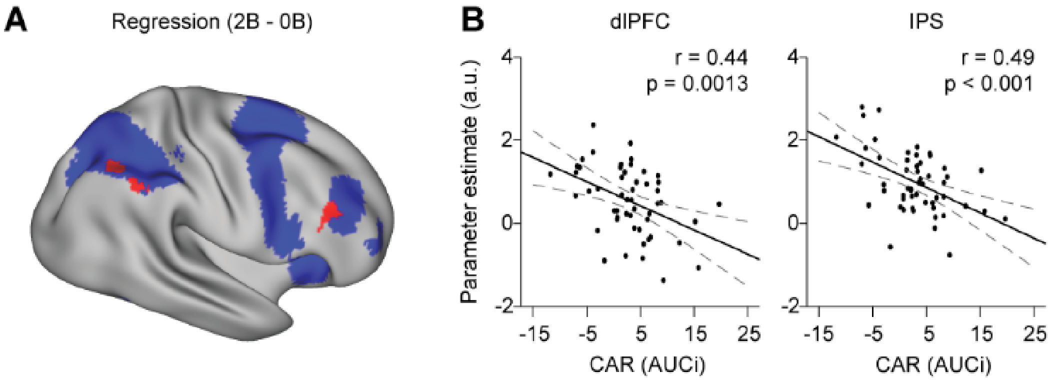


**Supplementary Fig. S2. Brain systems modulated by individual differences in CAR from Study 1. A**, A lateral view of significant clusters in the dlPFC and the IPS (in red), derived from a whole-brain regression analysis for WM-related neural activity (2- vs. 0-back) with CAR (AUCi) as the covariate of interest. Clusters in blue represent the main effect of WM. **B**, Scatter plots depict negative correlation of individual's CAR with dlPFC and IPS activity. Notes: R, right; 0B, 0-back; 2B, 2-back; CAR, cortisol awakening response; AUCi, area under the curve with respect to increase.

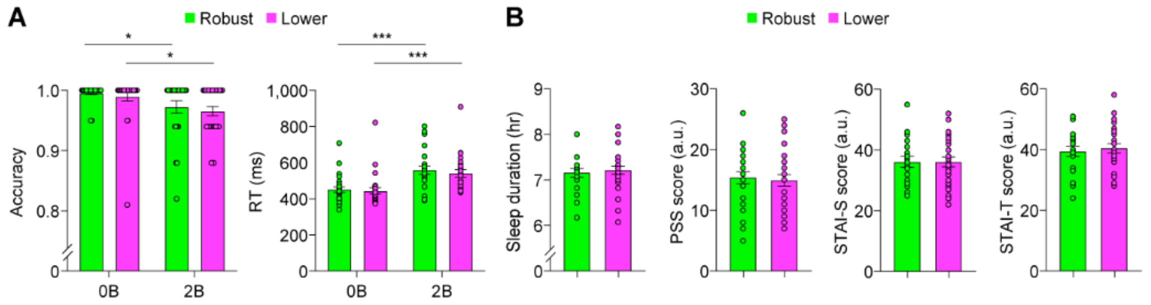


**Supplementary Fig. S3. Behavioural performance and psychological measurement from Study 1. A**, Bar graphs depict significant main effects of WM-loads for accuracy and reaction times (RTs). **B**, There was no significant group difference for sleep duration, PSS, STAI-S and STAI-T scores (*Table S1*). Error bars represent standard error of mean. Notes: 0B, 0-back; 2B, 2-back; PSS, perceived stress scale; STAI-S, state trait anxiety inventory-state; STAI-T, state trait anxiety inventory-trait; *, P < 0.05; ***, P < 0.001.

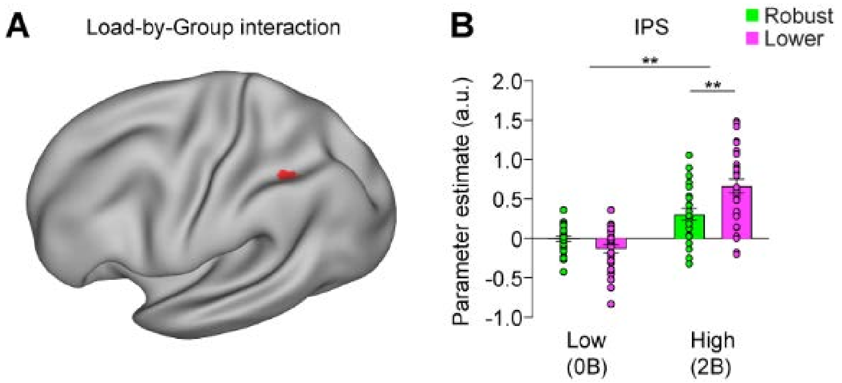


**Supplementary Fig. S4. Brain systems showing differential activation in individuals with robust- and lower-CAR from Study 1. A**, A lateral view of a significant cluster showing interaction effect between WM-load and Group in the left IPS. **B**, Bar graphs depict higher activation in the left IPS in individuals with lower- than robust-CAR only in high (2-back) but not low (0-back) task demand. Error bars represent standard error of mean. Notes: 0B, 0-back; 2B, 2-back; **, P < 0.01.

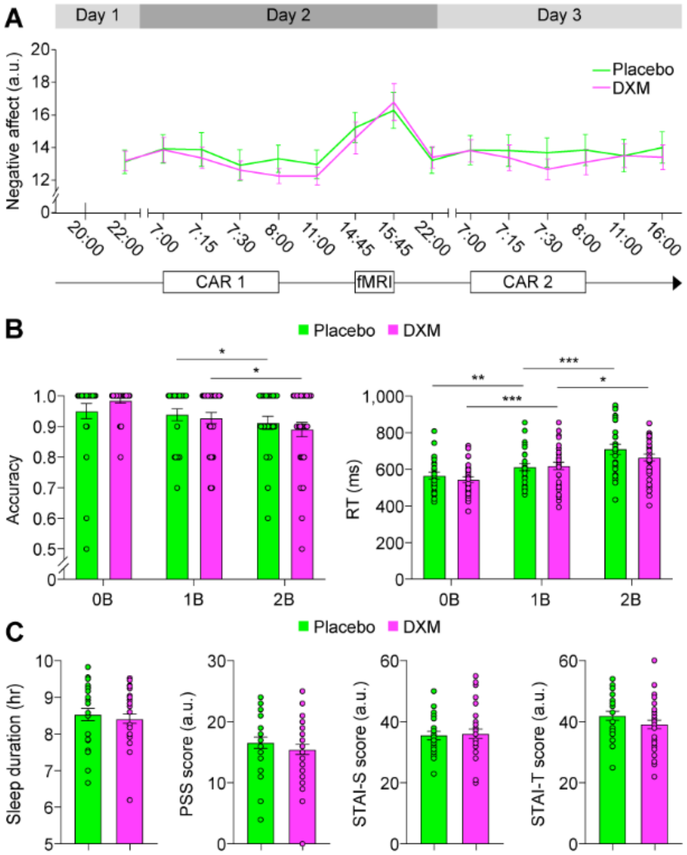


**Supplementary Fig. S5. Behavioural performance and psychological measurement from Study 2. A**, Subjective ratings of negative affect coinciding with 15 salivary samples, indicating no any side effect of DXM administration on participant's emotional well-being. **B**, Bar graphs depict significant main effects of WM-loads for both accuracy and RTs. **C**, Bar graphs depict no significant group difference for sleep duration, PSS, STAI-S and STAI-T scores (see *Table S1*). Error bars represent standard error of mean. Notes: 0B, 0-back; 2B, 2-back; PSS, perceived stress scale; STAI-S, state trait anxiety inventory-state; STAI-T, state trait anxiety inventory-trait; *, P < 0.05; **, P < 0.01; **, P < 0.001.

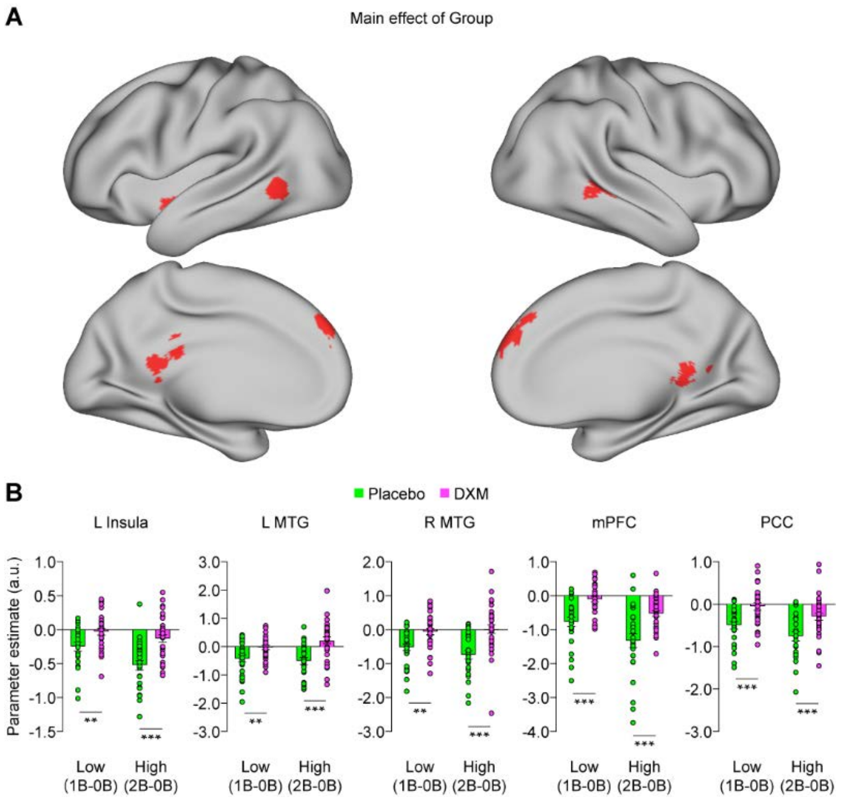


**Supplementary Fig. S6. Brain systems with differential activation in DXM and placebo groups from Study 2. A**, Lateral and medial views of significant clusters showing main effects of groups (i.e., DXM vs. placebo) in the insula, the middle temporal gyrus (MTG), the dorsal medial prefrontal cortex (dmPFC) and PCC. **B**, Bar graphs depict general hyper-activation in the insula, the MTG, the dmPFC and the PCC in DXM (vs. placebo) group regardless of WM loads. Error bars represent standard error of mean. Notes: L, left; R, right; 0B, 0-back; 1B, 1-back; 2B, 2-back; **, P < 0.01; ***, P < 0.001.

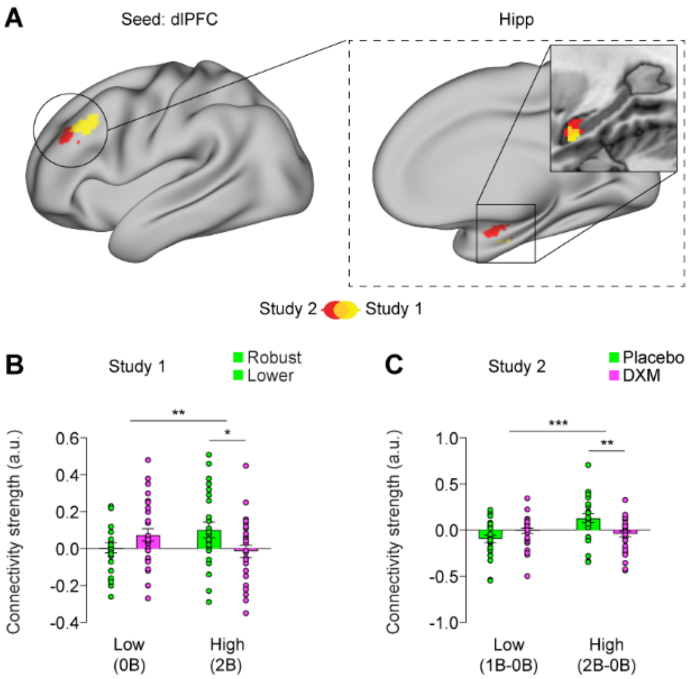


**Supplementary Fig. S7. Decreased prefrontal functional connectivity with the hippocampus in individuals with lower- and suppressed-CAR from Study 1 and 2. A**, Left: A lateral view of the dorsolateral PFC (in green and magenta for Study 1 and 2, respectively) as the seed for task-dependent functional connectivity, by using a generalized format of psychophysiological interaction (gPPI) approach. Right: A medial view and sagittal slice of a significant cluster in the left hippocampus showing an interaction between WM and groups. **B**, Bar graphs depict weaker functional connectivity of the dorsolateral PFC with the left hippocampus in individuals with lower- than robust-CAR only during high but not low task demand. **C**, Bar graphs depict weaker functional connectivity of the dorsolateral PFC with the hippocampus in DXM than placebo group only during high but not low task demand. Error bars represent standard error of mean. Notes: L, left; R, right; 0B, 0-back; 1B, 1-back; 2B, 2-back; *, P < 0.05; **, P < 0.01; ***, P < 0.001; n.s., not significant.

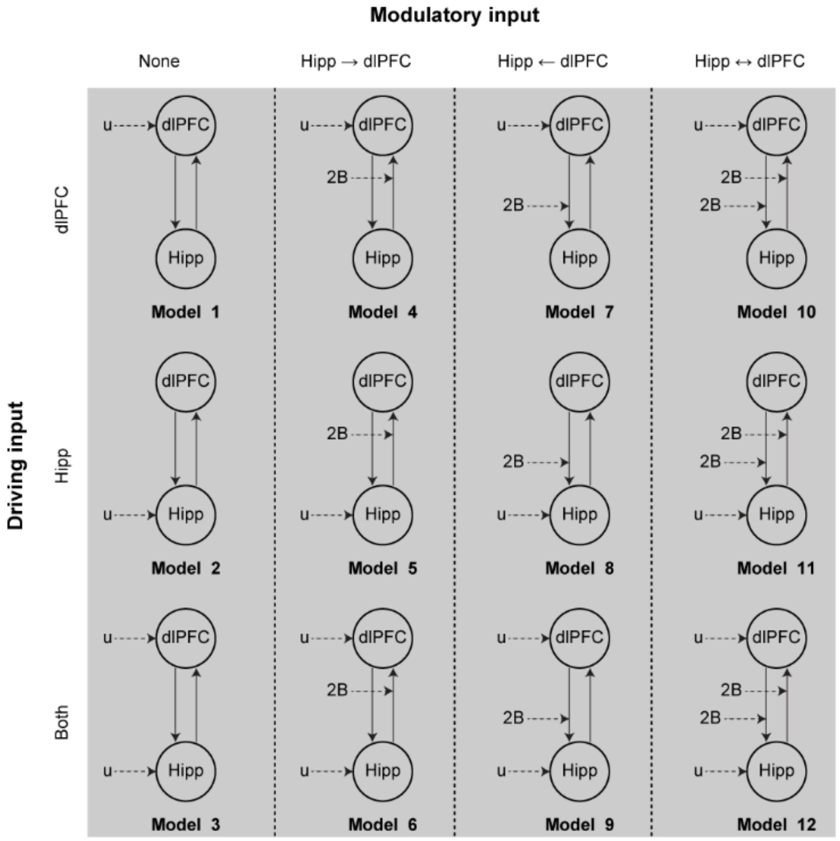


**Supplementary Fig. S8. Model specification in dynamic causal modeling (DCM).** For illustration purpose, the above 12 models included only 2-back condition as modulatory input. These models varied in the location of the driving input (i.e., either via the hippocampus, dlPFC, or both nodes), and the connections that could be modulated during 2-back conditions. Notes: dlPFC, dorsolateral prefrontal; Hipp, hippocampus; 2B, 2-back; u, driving input with all experimental conditions.

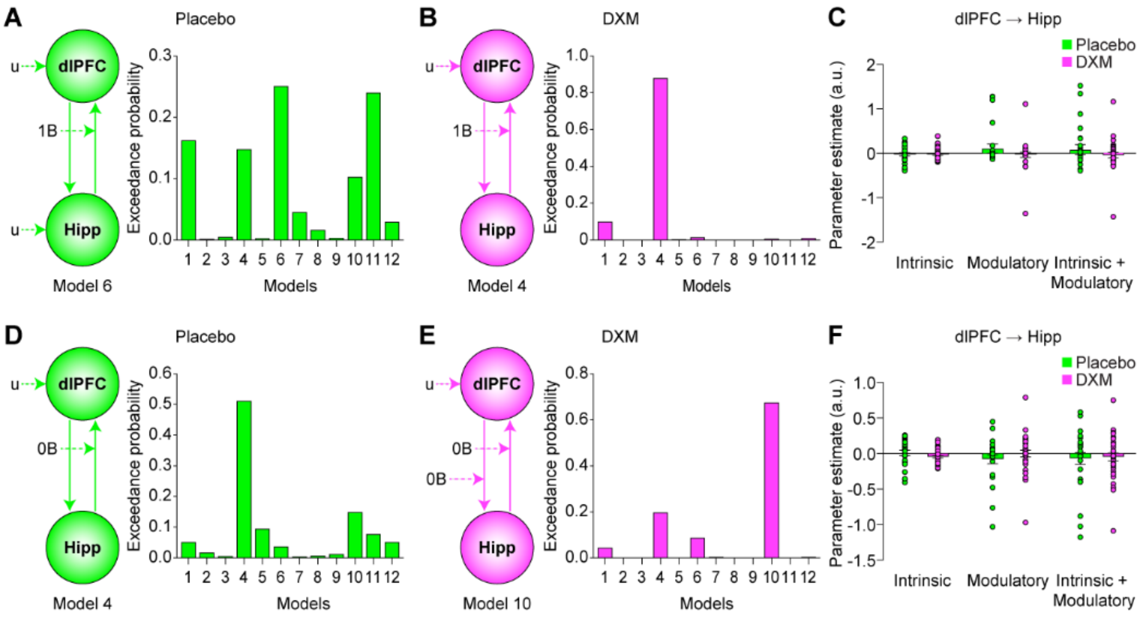


**Supplementary Fig. S9. Separated model selection within models with 1- and 0-back as modulatory in DCM analysis for DXM and placebo groups from Study 2. A**, For placebo, model evidence favored the 6th model with inputs to both the dlPFC and hippocampus driving the network, and with 1-back task only modulating dynamic influence from the hippocampus and dlPFC. **B**, For DXM, model evidence favored the 4th model with inputs to only the dlPFC driving the network, and with 1-back task only modulating dynamic influence from the hippocampus to dlPFC. **C**, Bar graphs depict no group difference in intrinsic, modulatory, and dynamic influence (i.e., intrinsic + modulatory) from the hippocampus to the dlPFC. **D**, For placebo, model evidence favored the 4th model with inputs to the dlPFC driving the network, and with 0-back task only modulating dynamic influence from the hippocampus and dlPFC. **E**, For DXM, model evidence favored the 10th model with inputs to only the dlPFC driving the network, and with 0-back task modulating dynamic influence of hippocampus with dlPFC bidirectionally. **F**, Bar graphs depict no group difference in intrinsic, modulatory and dynamic influence from the hippocampus to the dlPFC. Notes: dlPFC, dorsolateral prefrontal cortex; Hipp, hippocampus; 0B, 0-back; 1B, 1-back; 2B, 2-back; u, driving input with all experimental conditions.

### Supplementary Tables S1-S5

#### Supplementary Table S1. Participant demographics and psychological measurements

| | Study 1 | | | Study 2 | | |
|---|---|---|---|---|---|---|
| | Robust | Lower | P | Placebo | DXM | P |
| N | 28 | 24 | | 26 | 33 | |
| Age | 21.63±0.65 | 21.54±0.79 | 0.66 | 23.00±1.65 | 22.88±2.10 | 0.81 |
| (Range) | (21-23) | (20-23) | | (18-26) | (18-27) | |
| Sleep duration | 7.09±0.54 | 7.12±0.69 | 0.88 | 8.53±0.86 | 8.41±0.74 | 0.58 |
| PSS | 15.42±4.80 | 14.96±4.79 | 0.74 | 16.58±4.86 | 15.45±4.88 | 0.38 |
| STAI_S | 36.13±8.71 | 36.11±8.31 | 0.99 | 35.54±7.09 | 36.06±8.79 | 0.81 |
| STAI_T | 39.50±7.79 | 40.50±7.85 | 0.67 | 41.96±7.37 | 39.13±8.23 | 0.18 |

Mean (± standard deviation) of demographics and psychological measurements for Study (left) 1 and Study 2 (right). P values represent the significance of comparisons between groups. Notes: N, participants numbers; PSS, perceived stress scale; STAI-S, state trait anxiety inventory-state; STAI-T, state trait anxiety inventory-trait.

**Supplementary Table S2. Brain regions modulated by CAR from regression analyses (Study 1)**

| Region | L/R | BA | MNI (x, y, z) | | | *T* |
|---|---|---|---|---|---|---|
| **Regression: (2B and 0B) vs. Fixation** | | | | | | |
| Lingual gyrus | L | 19,37 | -24 | -54 | -10 | 4.45 |
| Thalamus | L | - | -22 | -16 | 4 | 4.34 |
| Middle cingulate cortex | L | 32,24 | -8 | 14 | 36 | 4.13 |
| Hippocampus | R | 35,28 | 22 | -24 | -16 | -3.85 |
| | | | 38 | -22 | -12 | -3.27 |
| **Regression: 2B vs. 0B** | | | | | | |
| Dorsal lateral prefrontal cortex | L | 9,8 | -40 | 24 | 44 | 3.79 |
| Intraparietal sulcus | R | 40 | 46 | -48 | 36 | 3.65 |
| | | | 52 | -46 | 46 | 3.44 |
| | | | 40 | -54 | 44 | 3.35 |
| Dorsal lateral prefrontal cortex | R | 46 | 48 | 30 | 24 | 3.37 |

Only clusters, significant at a height threshold of $P < 0.001$ and an extent threshold of $P < 0.05$ corrected on the whole brain level, are reported with local maxima in Montreal Neurological Institute (MNI) space. Clusters in the frontoparietal and hippocampal regions with prior hypotheses are in bold. Notes: L, left hemisphere; R, right hemisphere; BA, Brodmann’s area.

**Supplementary Table S3. Brain regions modulated by the CAR from ANOVA analyses (Study 1)**

| Region | L/R | BA | MNI (x, y, z) | | | *F* |
|---|---|---|---|---|---|---|
| **Main effect of Group** | | | | | | |
| Hippocampus | R | 28 | 28 | -20 | -16 | 19.34 |
| | | | 22 | -16 | -22 | 12.61 |
| **Group-by-Load interaction** | | | | | | |
| Inferior Parietal Lobule | L | 40 | -38 | -48 | 32 | 19.98 |
| Dorsal lateral prefrontal cortex | L | 9,8 | -44 | 24 | 38 | 16.95 |

Only clusters, significant at a height threshold of $P < 0.001$ and an extent threshold of $P < 0.05$ corrected on the whole brain level. Notes are the same as in Table S1.

**Supplementary Table S4. Brain regions modulated by the CAR from ANOVA analyses (Study 2)**

| Region | L/R | BA | *F* | MNI (x, y, z) | | |
|---|---|---|---|---|---|---|
| **Main effect of Group** | | | | | | |
| Medial prefrontal cortex | L | 9,6,10,8 | 22.95 | 2 | 56 | 36 |
| | | | 19.40 | -4 | 50 | 40 |
| | | | 13.55 | 14 | 56 | 32 |
| Cerebellum | L | - | 17.33 | -30 | -74 | -36 |
| | | | 15.14 | -28 | -84 | -32 |
| Insula | L | - | 16.74 | -34 | -10 | -6 |
| | | | 14.94 | -36 | -2 | -8 |
| Middle temporal gyrus | L | 37,21,39 | 16.35 | -56 | -56 | -2 |
| | | | 15.38 | 52 | -54 | 12 |
| Middle temporal gyrus | R | 21,22 | 15.71 | 66 | -34 | 0 |
| | | | 15.37 | 54 | -40 | -2 |
| | | | 14.48 | 42 | -52 | 14 |
| Hippocampus | R | 28 | 15.83 | 34 | -24 | -12 |
| **Group-by-Load interaction** | | | | | | |
| Dorsal lateral prefrontal cortex | L | 9 | 12.18 | -38 | 30 | 28 |

Only clusters, significant at a height threshold of $P < 0.001$ and an extent threshold of $P < 0.05$ corrected on the whole brain level. Notes are the same as in Table S1.

**Supplementary Table S5. Brain regions whose functional connectivity with the left dlPFC seed are modulated by the CAR from ANOVA analyses (Study 1 & Study 2)**

| Region | L/R | BA | *F* | MNI (x, y, z) | | |
|---|---|---|---|---|---|---|
| **Group-by-Load interaction (Study1)** | | | | | | |
| Posterior cingulate cortex | L&R | 31,23,30,29,7 | 14.49 | 0 | -50 | 26 |
| Postcentral Gyrus | R | 2,3,4,40,13 | 28.82 | 44 | -24 | 40 |
| Middle cingulate cortex | L&R | 31,6 | 15.17 | 10 | -28 | 42 |
| Medial Frontal Gyrus | R | 6,24,32 | 13.77 | 4 | 0 | 52 |
| **Hippocampus** | **L** | **-** | **10.76** | **-30** | **-16** | **-20** |
| **Hippocampus** | **R** | - | **11.00** | **40** | **-20** | **-20** |
| **Group-by-Load interaction (Study2)** | | | | | | |
| Lingual gyrus | L | 29,27 | 19.48 | -12 | -46 | 2 |
| | | | 13.63 | -12 | -38 | -2 |
| | | | 14.08 | 30 | 0 | 42 |
| Middle cingulate cortex | R | 24,31 | 17.78 | 14 | -12 | 44 |
| **Hippocampus** | **L** | **-** | **16.50** | **-30** | **-22** | **-16** |
| **Hippocampus** | **R** | **-** | **14.41** | **32** | **-10** | **-16** |

Only clusters, significant at a height threshold of $P < 0.001$ and an extent threshold of $P < 0.05$ corrected on the whole brain level. Given our priori hypothesis in the hippocampus from Study 1, we used a small volume correction procedure using the anatomically defined hippocampal mask. Notes are the same as in Table S1.
